# Risk of vascular events or death in different manifestations of cerebral small vessel disease: a 2-year follow-up study with a control group

**DOI:** 10.1101/151498

**Authors:** Jacek Staszewski, Renata Piusińska-Macoch, Bogdan Brodacki, Ewa Skrobowska, Katarzyna Macek, Adam Stępień

## Abstract

**Background and Purpose:** Natural course of cerebral small vessel disease (CSVD) has not yet been thoroughly studied. The aim of the single center study was to establish risk of vascular events or death in different manifestations of CSVD.

**Methods:** 150 consecutive, functionally independent patients with marked MRI features of CSVD and with recent lacunar stroke (n=52, LS), 20 with deep hemorrhagic stroke (HS), 28 with vascular parkinsonism (VaP), 50 with vascular dementia (VaD) and 55 controls (CG) with high atherothrombotic risk free of cerebrovascular events were prospectively recruited and followed for 24 months.

**Results:** Mean age and sex distribution were similar in CSVD and CG but patients with CSVD were less likely to have CAD (19% vs 40%, p=0,02) and tended to have higher prevalence of diabetes (54% vs 37%, p=0,11). The risk of vascular events or death was increased in any patients with moderate to severe white matter lesions at baseline MRI (HR 2,0; 95%CI 0,85-7,2), in CSVD (4,56; 95%CI 1,3-14,9) vs CG, regardless of its clinical manifestation: LS or HS (HR 4,70; 95%CI 1,3-16,2) and VaD or VaP (HR 4,59; 95%CI 1,3-15,7).

**Conclusions:** Patients with symptomatic CSVD regardless of the clinical (acute or chronic) manifestation had more than fourfold the risk of vascular events or death in 24 months of observation compared with controls with high atherothrombotic risk free of cerebrovascular events.

## Introduction

Cerebral small vessel disease (CSVD) is one of the most important and common vascular diseases of the brain caused by lacunar infarcts, white matter lesions (WMLs), microbleeds and intraparenchymal hemorrhage in the cerebral white and deep gray matter [1]. Although sharing similar risk factors, the pathophysiology of the disease is regarded as independent of that of atherosclerotic large artery disease. CSVD can cause several different types of distinct or overlapping clinical presentations, including recurrent lacunar strokes (LS), deep haemorrhagic strokes (HS), vascular dementia (VaD) and parkinsonism (VaP). Despite the radiological similarity, the short- and long-term prognosis of CSVD, especially HS or chronic symptomatic CSVD, are not well known. The risk of vascular events in acute and chronic manifestations of CSVD in comparison to atherothrombotic risk factors matched control group has not been adequately studied so far [2].

## Aim

In this single-center case-control study, we prospectively evaluated the short- and long-term prognosis in patients with non-disabling different clinical manifestations of CSVD and in control group followed for 24 months.

## Materials and Methods

The present investigation is embedded in the SHEF-CSVD Study (Significance of HEmodynamic and hemostatic Factors in the course of different manifestations of Cerebral Small Vessel Disease) [3]. The study group consisted of 150 consecutive patients with symptomatic CSVD and 55 controls (CG) without cerebrovascular disease but with high atherothrombotic risk. The patients were recruited from General or Neurological Outpatient Department and prospectively enrolled to the study between December 2011 and June 2014. The study protocol and methods have been thoroughly described elsewhere [3].

In brief, CSVD group consisted of consecutive patients with evidence of CSVD findings on neuroimaging (MRI), who were physically independent (modified Rankin Scale<3 and NIHSS<10 points) and had no severe dementia (MMSE≥12 points). The patients were diagnosed according to typical radiological and clinical picture, after exclusion of other neurodegenerative conditions with the use of clinical tools easily applied in clinical practice: LS-according to the OCSP Criteria, HS –parenchymal hemorrhage after vascular malformations, coagulopathies, trauma have been excluded; chronic VaP and VaD – after exclusion of other neurodegenerative conditions with the use of clinical tools easily applied in clinical practice: Hurtig criteria and NINDS-AIREN criteria with Modified Hachinski Ischemic Scale≥ 7 points, respectively [4, 5, 6]. Patients with post-stroke dementia or strategic single-infarct dementia were excluded. The date of the first-time recording of the VaP and VaD was used to establish the exact start date of these diseases. Basing on the nature of qualifying events, patients were further classified as acute CSVD (LS and HS) or chronic (VaD and VaP). Subjects without history of cerebrovascular disease and with normal MRI formed control group (CG). Participants from both groups were aged between 60 and 90 years. Patients with significant stenosis (>50%) of a major extracranial or intracranial artery, atrial fibrillation, non-CSVD related WMLs (e.g. due to migraine, vasculitis, multiple sclerosis, CADASIL, life expectancy of less than 6 months, and MRI contraindications were excluded.

### Procedure

Detailed neurological examination and assessments of cognitive function (mini-mental state examination (MMSE), disability (Barthel index, BI) and parkinsonism disease stage (Hoehn-Yahr) were performed at baseline and after 24-months. Patients were regarded dependent if gained ≤80 points in total BI scores [7]. Based on medical records, physical examination and comprehensive history available at baseline, we evaluated atherothrombotic risk factors, including tobacco use, diabetes, hyperlipidemia, hypertension, coronary artery disease (CAD), peripheral vascular disease and CKD. All patients received optimal medical treatment, according to guidelines. Data regarding vascular events during the study flow and/or cause of death (classified according to the ICD-10) was obtained both from treating physician, and/or general practitioner or medical records. Patients with incomplete follow-up were censored at the last time observation.

For MRI evaluation we used imaging with standard T2-weighted, FLAIR and gradient echo sequences. Periventricular and deep WMLs were visualized on T2 and PD/FLAIR images as ill-defined hyperintensities ≥5 mm. The simple modified Fazekas rating scale was used to estimate the extent of the per ventricular and deep WMLs [8]. Mild WMLs (grade 1) was defined as punctate lesions in the deep white matter with a maximum diameter of 9 mm for a single lesion and of 20 mm for grouped lesions. Moderate WMLs (grade 2) were early confluent lesions of 10-20 mm single lesions and >20 mm grouped lesions in any diameter, and no more than connecting bridges between the individual lesions. Severe WMLs (grade 3) were single lesions or confluent areas of hyperintensity of ≥20 mm in any diameter. All patients in CSVD group had at least grade 1 WMLs, those in CG were included only following normal MRI. All MRI scans were obtained using the same scanner (GE Healthcare 1.5T scanner).

The main outcome event was time to any vascular event (stroke – hemorrhagic or ischemic, myocardial infarction or peripheral vascular intervention) or death. Vascular death included death resulting from an acute myocardial infarction (MI), sudden cardiac death, heart failure, death due to stroke or cardiovascular procedures (CV) or hemorrhage, and death due to other CV causes [9]. Non-cardiovascular death was defined as any death with a specific cause that is not thought to be CV in nature. Undetermined cause of death referred to a death not attributable to one of the above categories. Ischemic and hemorrhagic strokes were defined based on typical clinical features with focal cerebral, spinal, or retinal dysfunction associated with a relevant radiological findings on a repeat brain MRI scan. Myocardial infarction was defined as a clinical or pathologic event caused by myocardial ischemia in which there was evidence of myocardial injury or necrosis with a rise and/or fall of cardiac biomarkers, along with supportive evidence in the form of typical symptoms, suggestive electrocardiographic changes, or imaging evidence of new loss of viable myocardium or new regional wall motion abnormality. Peripheral vascular intervention was defined as a catheter-based or open surgical procedure designed to improve arterial or venous blood flow or otherwise modify or revise vascular conduits.

This study complied with the Declaration of Helsinki. All participants from both groups signed an informed consent form. This study was approved by the local Medical Ethics Committee and was supported by the Polish Ministry of Science and Higher Education as a research project of the Military Institute of Medicine (Warsaw, Poland, study number N N402 473840).

### Statistical analysis

Log-normal data were compared using paired t-tests, non-normal data were analyzed using nonparametric tests, the chi-square test was used for comparisons of categorical variables. The difference between baseline risk factors was compared between CSVD patients and CG. Kaplan–Meier method was used to estimate the event-free survival, study groups were compared using Cox regression. Adjusted hazard ratios (HRs) and 95% CIs were calculated with fixed entry of a pre-defined set of potential confounders (age, sex, vascular co-morbidity and risk factors) selected on the basis of clinical plausibility and previous literature reviews and measured at the baseline visit. Dead or dependent at 24-months follow-up was analysed using Cox logistic regression. We did not control for baseline WMLs because it may distort estimates of change in longitudinal studies. A probability value of p<0.05 was considered significant. All data are presented as mean±SD values. All analyses were performed using Statistica 12 software (StatSoft Inc, USA).

## Results

### Patient characteristics

Study comprised of 205 patients: 150 with CSVD: 52 with first ever recent LS, 20 with HS, 28 with VaP and 50 with subcortical VaD and 55 controls. All controls had high CVD risk: 35 patients (63%) had documented symptomatic large artery disease (CAD or PAD), 3 (5,5%) had diabetes and CKD, 2 (3,6%) had diabetes alone. Mean age and sex distribution were similar in CSVD and CG but patients with CSVD were less likely to have CAD and tended to have higher prevalence of diabetes (Table 1). The estimated mean time from disease onset to enrollment was 17±2,1 days in HS; 17,5±3,8 days in LS; 25,5±11 months in VaD and 27,5±14 months in VaP. Grade 2 (n=83; 55,3%) or 3 (n=45;30%) WMLs were present in majority of patients with CSVD (80,3%) on baseline MRI.

**Table 1.**
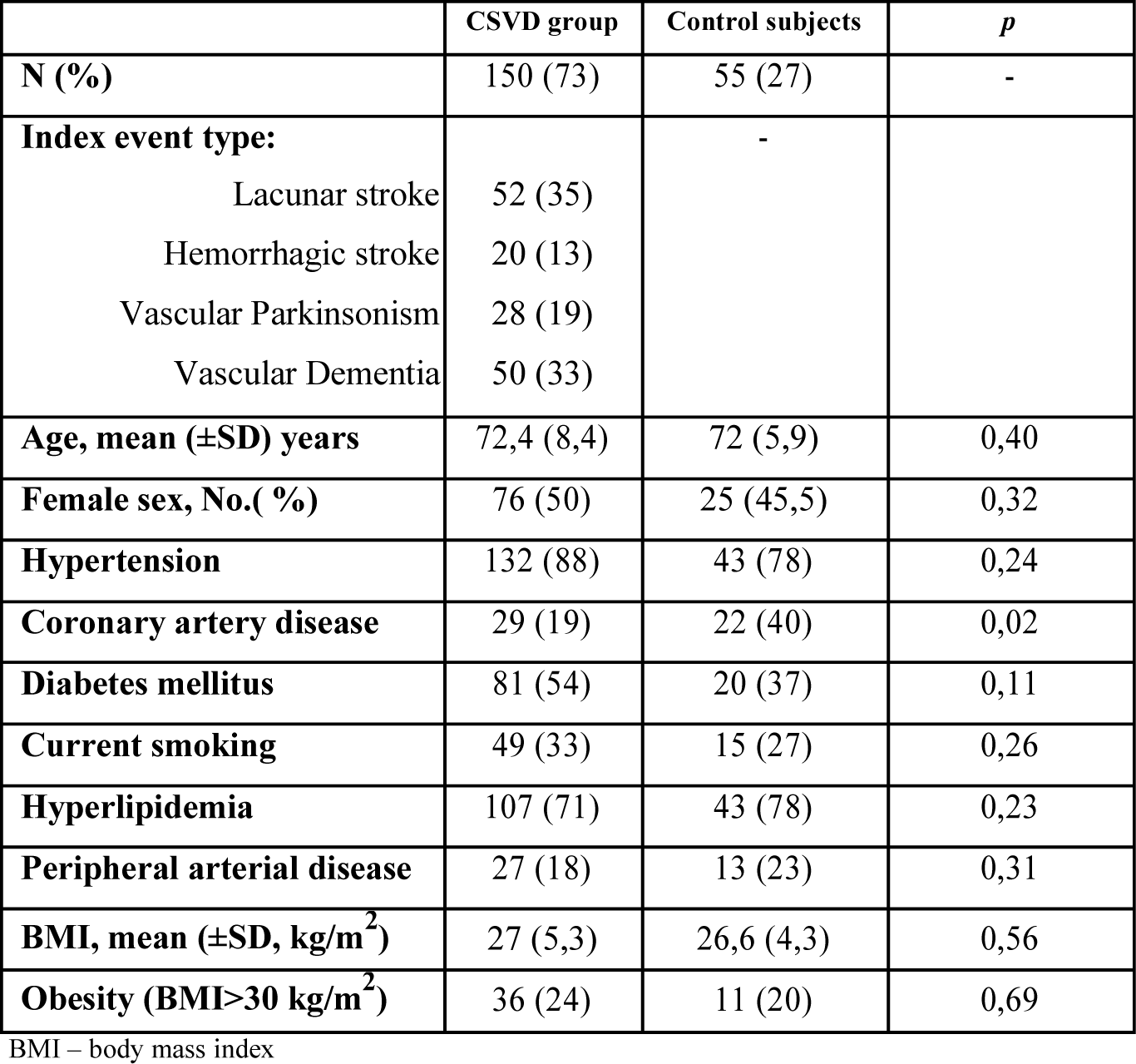
Baseline characteristics of patients with cerebral small vessel disease (CSVD) and control group (CG).

### All-cause mortality and vascular events

Information on the outcome was available from 97% of the enrolled patients, 4 patients (2 in VaD and 2 in CG) were lost to follow-up. The mean follow-up time in the studied population was 22,9±4,2 months (378,75 person-years): 22,4±4,8 months (271,25 person-years) in CSVD and 23,4±3 months (109,2 person-years) in CG (p=0,02). Eighteen patients (12%) died in CSVD group and none in CG group. The cause of death was vascular related in 9 (50%) patients (cardiac cause, n=8; hemorrhagic stroke, n=1), nonvascular in 3 (16%) (infection, n=2; malignant neoplasm, n=1) and undetermined in 6 (34%) patients. There were 30 nonfatal vascular events in CSVD and 3 events in CG (Table 2). The most frequent were ischemic strokes (IS) (n=25; all lacunar). Kaplan-Meier survival curves by SVD group showed significantly higher risk comparing with CG (log rank p=0,01) (Fig.1). Patients with CSVD were at a significantly higher risk (HR 4,56) for main outcome and IS (HR 9,73) compared with CG (Table 3). The sensitivity analyses for patients with chronic CSVD or Fazekas Grade 2 or 3 WMLs gave similar results. However, there were no significant differences between acute and chronic manifestations of CSVD with regard to risk of main outcome (HR 1,03; 95%CI 0,5-2,0) or IS (HR 0,57; 95%CI 0,2-1,4) but there was a trend toward increased risk of death from any cause in acute CSVD (HR 2,49; 95%CI 0,9-6,6) which was related to higher risk in patients with HS comparing with other CSVD groups (vs VaD; HR 11,1; 95%CI 2,9-41,8; LS HR 10,32; 95%CI 2,1-49,5 and VaP HR 2,97; 95%CI 0,8-10). In comparison to patients with mild WMLs patients with Fazekas Grade 2 or 3 had a trend toward a higher risk of vascular events od death (HR 2,0; 0,85-7,2) but not IS (HR 1,8; 0,4-8,7).

**Figure 1.**
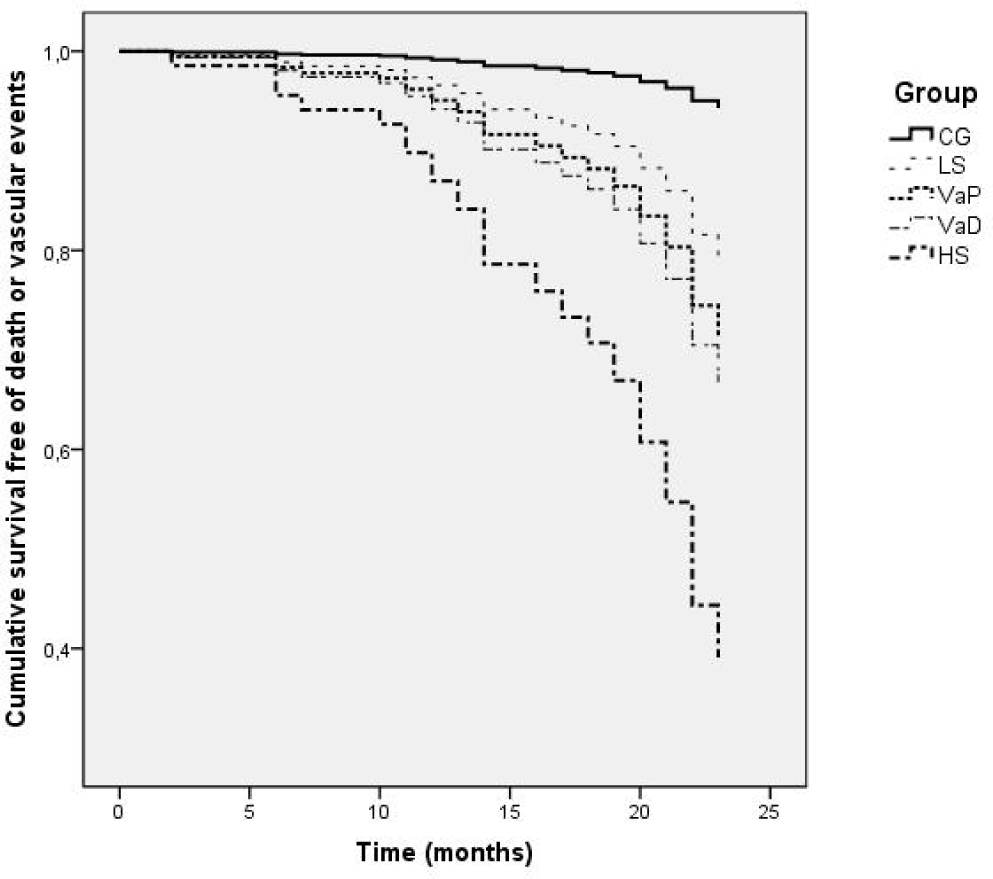
Kaplan-Meier curves for all-cause mortality and non fatal vascular events by grouped diagnostic category (log rank, p<0,01).

**Table 2.** The cause and numbers of different outcome events in patients with cerebral small vessel disease (CSVD)

| Studied group | Angioplasty for PAD | Ischemic stroke | Myocardial infarction | Intracranial hemorrhage | Vascular death | All-cause death | Main outcome |
| --- | --- | --- | --- | --- | --- | --- | --- |
| **CG, n(%)** | 0 | 1 (1,8) | 2 (3,6) | 0 | 0 | 0 | 3 (5,4) |
| **CSVD** | 2 (1,3) | 24 (16) | 3 (2) | 1 (0,7) | 9 (6) | 18 (12) | 39 (26) |
| **LS** | 2 (3,5) | 5 (9,6) | 2 (3,8) | 0 | 2 (3,8) | 2 (3,8) | 11 (21) |
| **HS** | 0 | 3 (15) | 0 | 1 (5) | 3 (15) | 9 (45) | 7 (35) |
| **VaP** | 0 | 2 (7,1) | 1 (3,5) | 0 | 1 (3,5) | 4 (14,2) | 4 (14,2) |
| **VaD** | 0 | 14 (28) | 0 | 0 | 3 (6) | 3 (6) | 17 (34) |

LS – lacunar stroke; HS – hemorrhagic stroke; VaP – vascular parkinsonism; VaD – vascular dementia ;PAD – peripheral arterial disease

**Table 3.** Sensitivity analyses in subsets of patients for main outcome, ischemic stroke and myocardial infarction.

|  | **n** | **Main outcome** | **Ischemic stroke** | **Myocardial infarction** |
| --- | --- | --- | --- | --- |
| **All patients** | 205 | 4,56 (1,3-14,9) | 9,73 (1,3-72,4) | 0,66 (0,1-3,9) |
| Excluding VaD, VaP | 127 | 4,70 (1,3-16,2) | 7,2 (0,89-57,8) | 1,00 (0,13-7,2) |
| Excluding HS and LS | 133 | 4,59 (1,3-15,7) | 12,6 (1,6-97,3) | 0,4 (0,3-4,5) |
| Excluding Grade 1 WMLs | 182 | 5,12 (1,5-16,8) | 10,9 (1,4-81,6) | 0,57 (0,8-4,1) |

VaP – vascular parkinsonism; VaD – vascular dementia; WMLs – white matter lesions
CSVD – cerebral small vessel disease; CG – control group
Values represent age and sex-adjusted HRs (95% CIs) for CSVD vs CG.

## Discussion

Despite this considerable clinical and societal impact, the causes and course in different clinical manifestations of CSVD are poorly understood. In this cohort study with 24 months follow-up, we have documented that CSVD increased more than fourfold the risk of future vascular events or death compared to patients with high atherothrombotic risk. The risk of death or vascular events was higher in CSVD and it did not depend on acute or chronic type of CSVD manifestations. This risk increased when moderate to severe WMLs were present at baseline MRI. During a two-year follow-up recurrent symptomatic strokes (both ischemic and hemorrhagic) were observed in 17% of patients with CSVD. Ischemic strokes were responsible for 61% of all vascular events in CSVD group, being the most frequent in patients presenting with VaD and HS. The long-term course of primary non-disabling hemorrhagic strokes and vascular dementia was in general unfavorable as 65% of patients with HS and 34% with VaD died or experienced recurrent vascular events during follow-up. We believe that, this risk in HS and LS groups was probably not linked with primary, qualifying events as the survival curves showed the highest incidence of events following the first year of observation. Exclusion of patients with atrial fibrillation and significant stenosis of extracranial or intracranial arteries in all patients resulted in a low incidence of events due to large vessel disease.

In comparison to strokes of different ethiologes, lacunar and hemorrhagic strokes in patients with CSVD were regarded in the past as less debilitating due to the smaller dimensions of brain lesions [10]. This view has been modified during the last decade when an excess of death and stroke recurrence has been consistently documented [11]. The risk of primary or secondary outcome was the highest following the first year of observation which was also well documented in the previous studies showing that the risk of death immediately after the onset of lacunar stroke was low but increased with time reaching 10-15% between 5 and 10 years and 60% of the patients died after 10 years [12]. In our cohort, the 2-year case-fatality rate in patients with LS was lower (3,8%) compared to previously reported data (8-15%) but the risk of recurrent stroke (9,6%) or other vascular events (7,3%) was similar to rates of 5-11% reported in prior prospective and epidemiological studies [13, 14]. These results were also similar to SPS3 trial in which, depending on the blood pressure control, patients with recent LS experienced 8% to 10% of recurrent strokes and 6,6%-7% died during 3,7 years of follow up [15]. It gave an annualized rate of recurrent stroke between 2,25% and 2,77% and death from all causes between 1,74%-1,8%. All recurrent IS in CSVD group in our study were of lacunar type, supporting a distinctive pathomechanism. The group of patients with HS was characterized by the highest risk of death and vascular events. Overall 24-month case fatality rate in this group was 45% what stood in concordance with previous reports in which at 3 months 18%-33,5% patients with nontraumatic, primary intracranial hemorrhage (ICH) died and less than half of them survived up to 1 year [16, 17]. In the metanalysis of the studies, the median 1-year mortality was 54.7% (range 46.0-63.6%) but these studies did not differentiate CSVD-related ICH from other causes [18]. Data pertaining to the risk of recurrent vascular events following ICH are scarce [19]. In our cohort, 20% of patients with index HS experienced recurrent strokes, and that risk was higher than previously reported in unselected ICH patients (12% at 3 years) but the number of cases in presented study was small [20]. More unfavorable course in our patients could be related to CSVD which is an independent predictor of worse outcomes in patients after spontaneous ICH [21]. We confirmed that the presence of severe WMLs is associated with a worse prognosis in terms of vascular events and death which is in concordance with the INTERACT2 study which showed that WMLs were linked with poor outcomes in acute ICH at 90 days [22]. Meta-analysis and systematic review of 46 prospective studies revealed the doubled risk of death among patients with WMLs in comparison to those without [23]. In this study we assessed the extent of WMLs solely, but there are other neuroimaging markers of cerebral CSVD, including cerebral microbleeds which represent poor prognostic markers of recurrent vascular events and functional recovery for stroke survivors [24, 25]. In a series of cohort studies based on the community-dwelling general population, the extent of WMLs increased the risk of IS, myocardial infarction, and mortality. The reason is unclear because it is still difficult to make a precise distinction between the direct effects of CSVD markers and bystander phenomena from shared vascular risk factors causing systemic micro- and macroangiopathies [26].

Our study documented that patients with lacunar or hemorrhagic strokes or VaD have a significantly higher risk of death and cardiovascular events compared with CG but on the other hand an excess in this risk was not observed in patients with VaP. Little is known about natural course of VaP and VaD caused by CSVD, which makes our data the more important. Although Parkinson’s disease has been consistently reported to be associated with a higher risk of all-cause mortality and stroke in various epidemiologic studies, previous studies of VaP were conducted on small groups probably due to imprecise diagnostic criteria [27]. In one prospective follow-up study of an unselected incident cohort of patients with VaP (n=38) and age-sex matched controls (n=262), patients with VaP had increased mortality (HRs 6.85) and 96% rate of death or dependency at 3 years [28]. In our study, the risk of vascular death and vascular events including stroke in VaP was almost three-fold higher than in CG but it was not statistically significant. As it was probably caused by a low number of subjects in that group, on the other hand the risk for primary and secondary outcomes in VaP did not significantly differ from patients with LS and VaD and the risk of secondary outcome was significantly higher than in CG.

We demonstrated significantly higher risk of vascular death and vascular events in VaD comparing to CG, with a rate of all cause death of 14,2%, which was consistent with the previous reports. The risk of death was even lower comparing to a study in which 24% individuals with VaD followed for 2 years died [29]. The higher risk of death and a greater risk of stroke and death from coronary events in that population probably resulted from an older age of population studied. Although VaD and VaP are not acute life-threatening disorders, our study documented a more unfavorable course in all clinical manifestations including chronic CSVD which demonstrated high risk of all-cause death and vascular events (HRs 4.59-5.40) compared to controls. An even more pronounced risk (relative risk 11.1) was reported previuosly when patients with different types of dementia were compared with controls, which may be, however attributed to their significantly older age (mean 85 years) than in current study [30]. The important cause of death in these patients compared to general population were dementia alone and cerebrovascular diseases. The excess in the risk of nonvascular death in this population can be explained. As these diseases progress patients become increasingly frail, which gives rise to complications like swallowing impairment, incontinence or falls. It is interesting that even in the present study an apparent similarity emerged between the course of acute and chronic CSVD what can further prove common pathogenetic mechanisms of these entities. Although an obviously different disease course in patients with CSVD and those with CG matched for similar risk factors suggest pathophysiology of CSVD may be independent from that of atherosclerotic large artery disease, we found no significant difference in risk of MI in patients with CSVD (2%) and CG (3,6%). Similar data was reported by Yamamoto Y et al with 3,3% patients experiencing MI following lacunar infarct [31]. That also suggests that “nonvascular” component and nonvascular contributing mechanisms may play a role in the cause and long-term course of CSVD [32].

The results from our study suggest that regardless of the clinical manifestation, the course of CSVD is generally poor. It would be important for clinicians to identify high risk patients, and especially those with chronic CSVD, to implement more effective preventive strategies. Precision in prediction of mortality can be improved f.e. by imaging modalities. Variability in clinical presentations of CSVD and unfavorable course may be attributable to either the poor control or the aftermath of traditional vascular risk factors. The association between CSVD and clinical outcome can depend upon several underlying mechanisms. Cerebral small vessel disease reflects potential fragility of the brain and an increased susceptibility to ischemic or hemorrhagic brain damage. These changes may increase the risk of not only overt infarction from vessel occlusion but also chronic ischemia from reductions in perfusion and gas transfer in the white matter and deep brain structures [1, 33]. It has been suggested that areas with extensive WMLs may be poorly supplied by collateral compensation, which may lead to IS extension, decreased neuronal connectivity and poor brain compensation and recovery [34].

Our study strengths include its prospective design and parallel 24 month observation of patients with well characterized different CSVD manifestations including rarely studied chronic VaP and VaD compared with control population sharing similar risk factors. Furthermore, CG was a random sample of the CSVD free population matched in respect to age, sex and vascular risk factors that attended the same Outpatient Department at approximately the same time as CSVD group, which is a design that minimizes the potential for selection bias and confounding. Another advantage is the complete follow-up for mortality, and single-center design which allowed us to consistently collect all measures. Our study has also some limitations. The major weakness is the small number of patients and controls included. The total number of outcome events, particularly recurrent strokes, was relatively small. Patients with VaP and VaD included in the present study were in an advanced stage of their disease therefore it remains unknown whether our results can be applied to less severely affected patient. The overall study period (24 months) could ideally have been longer to reach more significant results. Therefore, future studies are needed to develop a predictive risk score for chronic manifestations of CSVD, and to clarify the interaction between degenerative and vascular processes in their development. As it is impossible to achieve a 100-percent accuracy in intravital diagnosis of parkinsonism without pathological confirmation, there may be in our study, as well as in any previous ones, some imprecision in that matter despite the fact that we applied strict diagnostic criteria based on all available clinical data.

## Summary

The main outcome of our study is that, patients with CSVD have an excess risk of death and increased rate of lacunar stroke in comparison with controls with high atherothrombotic risk free of cerebrovascular disease. The study demonstrated that different clinical manifestations in CSVD, both acute or chronic have similar unfavorable prognosis in the 24 month of observation. The precise mechanisms by which CSVD mediates poor outcomes are unclear and require additional investigations. Further studies are warranted to evaluate the risk of CSVD in the long run.

